# Boolean Network Modeling Identifies Cognitive Resilience in the First Murine Model of Asymptomatic Alzheimer’s Disease

**DOI:** 10.1101/2025.06.11.659207

**Authors:** Suborno Jati, Sahar Taheri, Satadeepa Kal, Subhash C. Sinha, Brian P. Head, Sushil K. Mahata, Debashis Sahoo

**Author notes:** Senior corresponding authors: Debashis Sahoo, Ph.D.; Associate Professor, Department of Pediatrics, University of California San Diego; 9500 Gilman Drive, MC 0703, Israni Biomedical Research Facility Room No 2119; La Jolla, CA 92093-0703., Sushil K. Mahata, Ph.D.; Metabolic Physiology & Ultrastructural Biology Laboratory; Department of Medicine; University of California, San Diego (0732); 9500 Gilman Drive; La Jolla, CA 92093-0732. Equal contribution (Suborno Jati, Sahar Taheri).

## Abstract

Alzheimer’s disease (AD) is a progressive neurodegenerative disorder defined by amyloid beta (Aβ) plaques and neurofibrillary tangles (NFTs), yet approximately 20–30% of aged individuals exhibit these hallmark lesions without developing cognitive impairment—a clinically silent condition termed asymptomatic AD (AsymAD). The molecular basis of this cognitive resilience remains poorly understood due to a lack of mechanistic models. Here, we integrate systems-level Boolean network modeling with *in vivo* validation to define the transcriptomic logic of AsymAD and uncover a novel preclinical model. Using Boolean implication networks trained on large-scale human cortical RNA-seq datasets, we identified a robust and invariant AD gene signature that accurately stratifies disease states across independent datasets. Application of this signature to Chromogranin A– deficient PS19 mice (CgA-KO/PS19) revealed a unique resilience phenotype: male mice developed AD-like molecular and neuropathological profiles in the pre-frontal cortex yet retained intact learning and memory. Female CgA-KO/PS19 mice displayed even greater protection, including reduced Tau phosphorylation and preserved synaptic ultrastructure. These findings establish the first validated murine model of AsymAD and identify CgA as a modifiable node linking neuroendocrine signaling, Tauopathy, and cognitive preservation. This work provides a scalable platform to probe sex-specific resilience, uncover early-stage biomarkers, and accelerate preventive therapeutic development in AD.

## Introduction

Alzheimer’s disease (AD) is a progressive, fatal neurodegenerative disorder characterized by extracellular deposition of amyloid beta (Aβ) in neuritic plaques and intracellular accumulation of hyperphosphorylated Tau in neurofibrillary tangles (NFTs) ^1–4^. These hallmark pathologies have long been considered necessary correlates of clinical dementia. Yet, epidemiological and neuropathological studies reveal that ∼20–30% of older individuals harbor extensive amyloid and Tau pathology without exhibiting cognitive impairment prior to death ^5–7^. This clinically silent phase, termed asymptomatic Alzheimer’s disease (AsymAD), presents a critical but understudied window for early intervention and disease interception.

AsymAD is not simply a prodrome of overt AD; rather, it represents a unique neurobiological state defined by the coexistence of pathological burden and cognitive resilience. Human studies have shown selective neuronal hypertrophy in regions such as the anterior cingulate cortex and CA1 hippocampus ^8–10^, accompanied by a distinct immunomodulatory landscape—characterized by quiescent or anti-inflammatory glial phenotypes, reduced microglial activation, and suppression of pro-inflammatory cytokines ^11, 12^. However, the molecular programs that confer resilience in the face of Tau pathology remain poorly defined ^13–16^. This knowledge gap is partly due to the absence of suitable animal models that recapitulate the dissociation between pathology and cognition, and partly due to a lack of systems-level tools to decode the transcriptional logic of resilience.

Recent studies highlight the spatial and cellular heterogeneity of AD and suggest that local tissue environments—particularly glial and vascular niches surrounding plaques—modulate the trajectory of neurodegeneration ^16–18^. Understanding how these microenvironments are shaped in AsymAD requires computational frameworks that move beyond static gene lists to capture robust, invariant regulatory signatures. Boolean logic modeling offers one such approach, revealing gene-gene relationships that persist across individuals, disease stages, and datasets.

Chromogranin A (CgA), a secretory granule protein expressed in neuroendocrine and neuronal tissues ^19–21^, has emerged as a candidate modulator of AD vulnerability ^22,23–26^. CgA is elevated in the cerebrospinal fluid (CSF) of AD patients ^27, 28^, where its levels correlate with total and phosphorylated Tau, and it colocalizes with Tau aggregates in NFTs. Our recent work demonstrated that CgA deletion in PS19 Tauopathy mice (CgA-KO/PS19) reduces Tau burden, delays disease progression, and improves behavioral outcomes ^28^. These findings suggest that CgA may act upstream of Tau pathology— possibly via regulation of catecholaminergic tone, neuroinflammation, and glial reactivity.

In this study, we combine transcriptomic modeling with *in vivo* validation to establish a preclinical model of AsymAD. Using Boolean Network Explorer (BoNE) ^29^, we identified a core, invariant AD gene signature from human cortical RNA-seq datasets and applied it to CgA-KO/PS19 mice. We report that male CgA-deficient PS19 mice exhibit robust AD-like molecular profiles in the hippocampus and entorhinal cortex yet maintain intact cognitive performance. Notably, female mice show even greater resilience, with reduced Tau phosphorylation and preserved synaptic ultrastructure. These findings define the CgA-KO/PS19 mouse as the first experimentally validated model of AsymAD. By linking systems-level gene networks to sex-specific resilience phenotypes, this work provides a mechanistic foundation for early detection, biomarker discovery, and therapeutic targeting of preclinical AD.

## Methods

### Animals

All animal experiments were approved by the Institutional Animal Care and Use Committee (IACUC) at the University of California, San Diego (UCSD) and the VA San Diego Healthcare System, and were conducted in accordance with NIH guidelines and the ARRIVE reporting standards.

Chromogranin A knockout (CgA-KO) mice were originally generated on a mixed genetic background (50% 129/SvJ and 50% C57BL/6) using a *Cre*-*loxP* gene-targeting strategy to achieve congenital and whole-body deletion of the *Chga* gene ^30^. These CgA-KO mice were subsequently backcrossed to C57BL/6J mice for 7 generations to establish a congenic C57BL/6 background. To generate mice in the B6C3F1/J background, the C57BL/6J CgA-KO mice were backcrossed to B6C3F1/J mice for four generations. These mice were subsequently crossed with PS19 heterozygous mice (B6C3F1/J background; Jackson Laboratory, stock #008169) to generate *CgA-KO/PS19* experimental animals ^28^.

All mice were housed in a temperature- and humidity-controlled facility on a 12-hour light/12-hour dark cycle, with ad libitum access to food and water. Animals were fed a standard normal chow diet (NCD; 14% kcal from fat; LabDiet 5P00).

### Euthanasia and Tissue Collection

Mice were euthanized in accordance with IACUC protocols and the 2020 AVMA Guidelines for the Euthanasia of Animals. Animals were placed in an induction chamber pre-filled with 3–5% isoflurane in oxygen (flow rate: 1–2 L/min) and monitored for loss of the righting reflex and absence of response to toe pinch, indicating a surgical plane of anesthesia. While under deep anesthesia, tissues were harvested, and euthanasia was completed via exsanguination.

### Genotyping

Mice were ear-tagged, and tail biopsies were collected for genotyping. Genomic DNA was extracted using the AccuStart Genotyping Kit (QuantaBio), and PCR amplification was performed using the AccuStart GelTrack PCR SuperMix.

*PS19 Genotyping Primers:*

- Forward (WT and mutant): 5′-TTG AAG TTG GGT TAT CAA TTT GG-3′
- Reverse (WT): 5′-TTC TTG GAA CAC AAA CCA TTT C-3′
- Reverse (Mutant): 5′-AAA TTC CTC AGC AAC TGT GGT-3′

*Chga Genotyping Primers:*

- Forward: 5′-GTA GCA TGG CCA CTA CCC AG-3′
- Reverse: 5′-ATC CTT CAG AGC CCC TCC TT-3′

### Building a Comprehensive Database of Alzheimer’s disease Datasets

Publicly available microarray and RNASeq databases were downloaded from the National Center for Biotechnology Information (NCBI) Gene Expression Omnibus (GEO) website ^31–33^. Gene expression summarization was performed by normalizing Affymetrix platforms by RMA (Robust Multichip Average) ^34, 35^ and RNASeq platforms by computing TPM (Transcripts Per Millions) ^36^ values whenever normalized data were not available in GEO. We used log2(TPM+1) as the final gene expression value for analyses. We also used publicly data normalized using RPKM ^37^, FPKM ^38, 39^, TPM ^40, 41^, and CPM ^42, 43^. In the context of Affymetrix microarray data we believe that RMA works better than MAS 5.0 ^44^. List of training and validation datasets used were provided in Supplementary Data.

### StepMiner Analysis

StepMiner is a computational tool that identifies step-wise transitions in a time-series data ^45^. StepMiner performs an adaptive regression scheme to identify the best possible step up or down based on sum-of-square errors. The steps are placed between time points at the sharpest change between low expression and high expression levels, which gives insight into the timing of the gene expression-switching event. To fit a step function, the algorithm evaluates all possible step positions, and for each position, it computes the average of the values on both sides of the step for the constant segments. An adaptive regression scheme is used that chooses the step positions that minimize the square error with the fitted data. Finally, a regression test statistic is computed as follows:

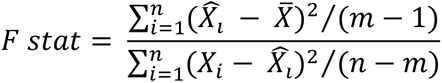

Where *X_i_* for *i* = 1 to *n* are the values, 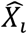 for *i* = 1 to *n* are fitted values. m is the degrees of freedom used for the adaptive regression analysis. 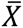 is the average of all the values: 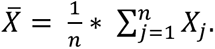 For a step position at k, the fitted values 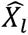 are computed by using 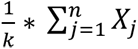 for *i* = 1 to *k* and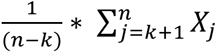 for *i* = *k* + 1 to *n*.

### Boolean Analysis

**Boolean logic** is a simple mathematical relationship of two values, i.e., high/low, 1/0, or positive/negative. The Boolean analysis of gene expression data requires the conversion of expression levels into two possible values. The ***StepMiner*** algorithm is reused to perform Boolean analysis of gene expression data ^29^. **The Boolean analysis** is a statistical approach which creates binary logical inferences that explain the relationships between phenomena. Boolean analysis is performed to determine the relationship between the expression levels of pairs of genes. The ***StepMiner*** algorithm is applied to gene expression levels to convert them into Boolean values (high and low). In this algorithm, first the expression values are sorted from low to high and a rising step function is fitted to the series to identify the threshold. Middle of the step is used as the StepMiner threshold. This threshold is used to convert gene expression values into Boolean values. A noise margin of 2-fold change is applied around the threshold to determine intermediate values, and these values are ignored during Boolean analysis. In a scatter plot, there are four possible quadrants based on Boolean values: (low, low), (low, high), (high, low), (high, high). A Boolean implication relationship is observed if any one of the four possible quadrants or two diagonally opposite quadrants are sparsely populated. Based on this rule, there are six kinds of Boolean implication relationships. Two of them are symmetric: equivalent (corresponding to the positively correlated genes), opposite (corresponding to the highly negatively correlated genes). Four of the Boolean relationships are asymmetric, and each corresponds to one sparse quadrant: (low => low), (high => low), (low => high), (high => high). BooleanNet statistics is used to assess the sparsity of a quadrant and the significance of the Boolean implication relationships ^29, 46^. Given a pair of genes A and B, four quadrants are identified by using the StepMiner thresholds on A and B by ignoring the Intermediate values defined by the noise margin of 2-fold change (+/- 0.5 around StepMiner threshold). Number of samples in each quadrant are defined as a_00_, a_01_, a_10_, and a_11_ which is different from X in the previous equation of F stat. Total number of samples where gene expression values for A and B are low is computed using the following equations.

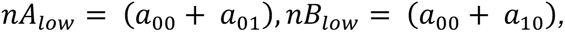

Total number of samples considered is computed using following equation.

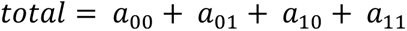

Expected number of samples in each quadrant is computed by assuming independence between A and B. For example, expected number of samples in the bottom left quadrant 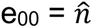 is computed as probability of A low ((a_00_ + a_01_)/total) multiplied by probability of B low ((a_00_ + a_10_)/total) multiplied by total number of samples. Following equation is used to compute the expected number of samples.

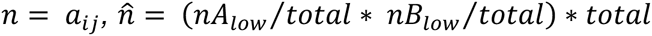

To check whether a quadrant is sparse, a statistical test for (e_00_ > a_00_) or 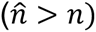 is performed by computing S_00_ and p_00_ using following equations. A quadrant is considered sparse if S_00_ is high 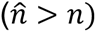 and p_00_ is small.

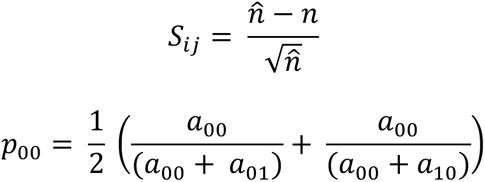

A suitable threshold is chosen for S_00_ > sThr and p_00_ < pThr to check sparse quadrant. A Boolean implication relationship is identified when a sparse quadrant is discovered using following equation.

***Boolean Implication*** = (*S_ij_* > sThr, *p_ij_* < pThr)

A relationship is called Boolean equivalent if top-left and bottom-right quadrants are sparse.

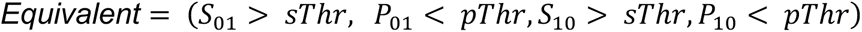

Boolean opposite relationships have sparse top-right (a_11_) and bottom-left (a_00_) quadrants.

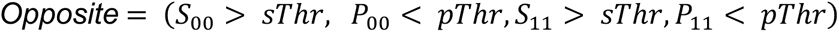

Boolean equivalent and opposite are symmetric relationship because the relationship from A to B is same as from B to A. Asymmetric relationship forms when there is only one quadrant sparse (A low => B low: top-left; A low => B high: bottom-left; A high=> B high: bottom-right; A high => B low: top-right). These relationships are asymmetric because the relationship from A to B is different from B to A. For example, A low => B low and B low => A low are two different relationships.

A low => B high is discovered if the bottom-left (a_00_) quadrant is sparse and this relationship satisfies following conditions.

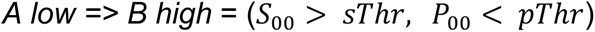

Similarly, A low => B low is identified if the top-left (a_01_) quadrant is sparse.

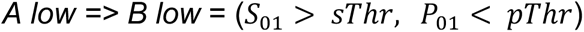

A high => B high Boolean implication is established if the bottom-right (a_10_) quadrant is sparse as described below.

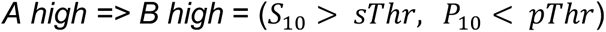

Boolean implication A high => B low is found if the top-right (a_11_) quadrant is sparse using following equation.

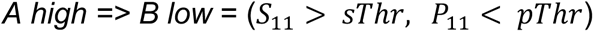

For each quadrant a statistic S_ij_ and an error rate p_ij_ is computed. S_ij_ > sThr and p_ij_ < pThr are the thresholds used on the BooleanNet statistics to identify Boolean implication relationships.

Boolean analyses in the test dataset GSE125583 uses a threshold of sThr = 3 and pThr = 0.1. These thresholds are exactly same as the previously used thresholds sThr = 3 and pThr = 0.1 for BooleanNet ^44, 45, 47^. False discovery rate is computed for these thresholds (FDR < 0.00001) by using randomly permuting gene expression data in GSE125583.

### Boolean Network Explorer (BoNE)

Boolean network explorer (*BoNE*) provides an integrated platform for the construction, visualization and querying of a network of progressive changes underlying a disease or a biological process in three steps (**Fig. 1a**): First, the expression levels of all genes in these datasets were converted to binary values (high or low) using the StepMiner algorithm. Second, gene expression relationships between pairs of genes were classified into one-of-six possible Boolean Implication Relationships (BIRs), two symmetric and four asymmetric, and expressed as Boolean implication statements. This offers a distinct advantage from conventional computational methods (Bayesian, Differential, etc.) that rely exclusively on symmetric linear relationships in networks. The other advantage of using BIRs is that they are robust to the noise of sample heterogeneity (i.e., healthy, diseased, genotypic, phenotypic, ethnic, interventions, disease severity) and every sample follows the same mathematical equation, and hence is likely to be reproducible in independent validation datasets. Third, genes with similar expression architectures, determined by sharing at least half of the equivalences among gene pairs, were grouped into clusters and organized into a network by determining the overwhelming Boolean relationships observed between any two clusters. In the resultant Boolean implication network, clusters of genes are the nodes, and the BIR between the clusters are the directed edges; *BoNE* enables their discovery in an unsupervised way while remaining agnostic to the sample type.

**Figure 1:**
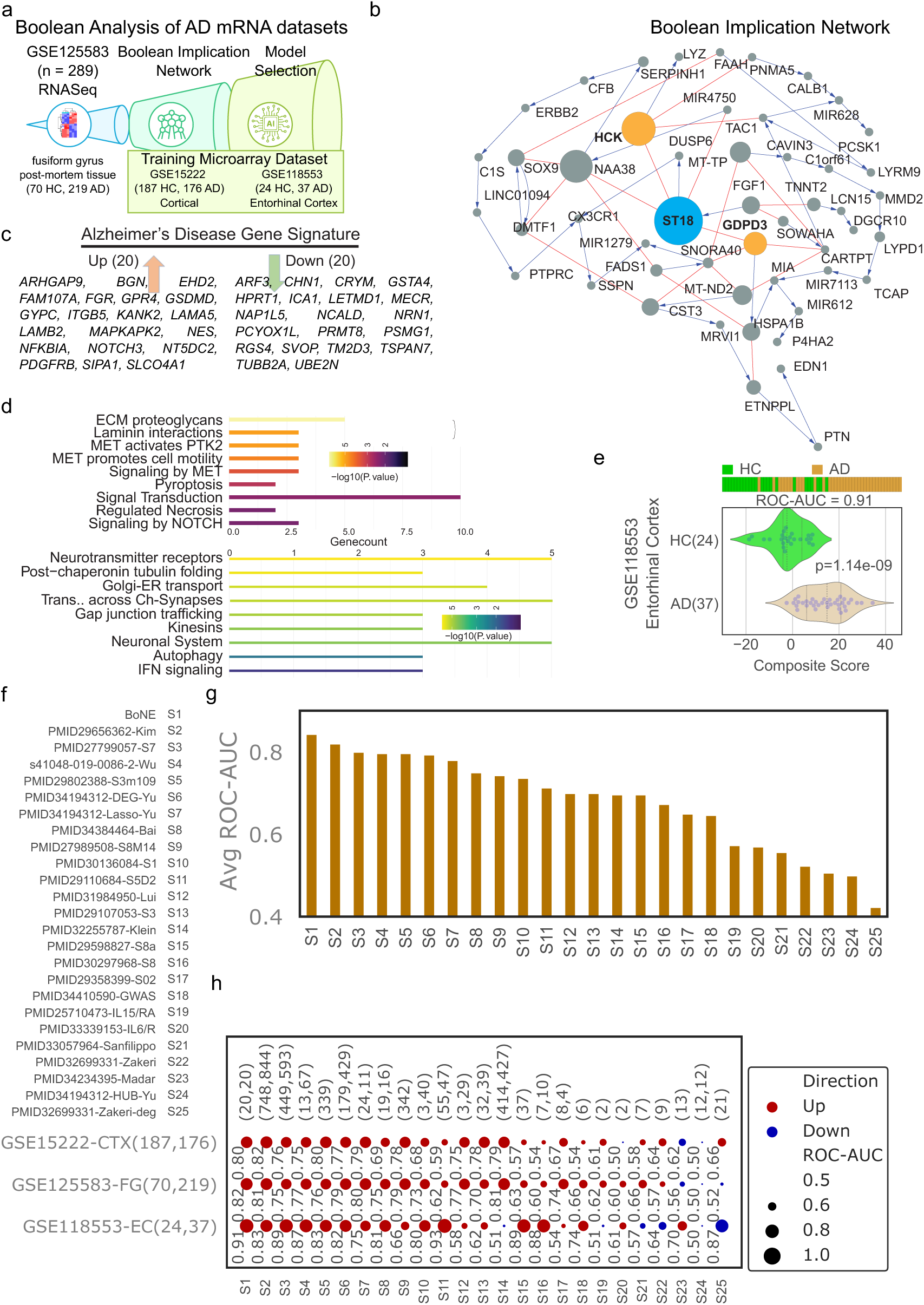
Boolean Analysis of Alzheimer’s disease mRNA datasets. (**a**) Schematic representation of the Boolean analysis for Alzheimer’s disease (AD) datasets. (**b**) Network of clusters with representative gene name (size of circle = number of genes in the cluster) and Boolean implication relationships (red = hilo, blue – lolo). Top three clusters highlighted include differentially expressed genes between healthy control (HC) and AD brain samples. (**c**) AI/ML predicted gene signatures to distinguish HC and AD samples based on the network in panel b and three training datasets (GSE125583 Fusiform Gyrus, GSE15222 Cortical, GSE118553 Entorhinal Cortex). (**d**) Reactome analysis of the up/down regulated genes. (**e**) The composite score of the Boolean AD models was evaluated in GSE118553 Entorhinal Cortex, ordering of samples were visualized using bar plot and its distribution in each category is shown in violin plots. The ROC-AUC value and p-value from a two-sided unpaired T-test with unequal variance were computed to assess the Boolean model’s ability to distinguish between 24 HC (colored green) and 37 AD samples (colored orange) from the Entorhinal Cortex (GSE118553). (**f**) 24 publicly available gene signatures are ranked and compared to the trained Boolean AD model based on their average ROC-AUC values in distinguishing healthy control (HC) and Alzheimer’s disease (AD) samples across the three training datasets. (**g**) The results are presented in a bar plot, with the x-axis representing the ranked gene signatures and the average ROC-AUC values ordered from highest to lowest. (**h**) ROC-AUC analysis in bubble plots represent composite gene signature analyses of 24 previously published gene signatures and Boolean AD model (S1) in the three different training datasets. Up regulated gene signatures are colored red and down regulated gene signatures are colored blue. Bubble size corresponds to ROC-AUC, significance of p values are shown as *** (p < 0.001), ** (p < 0.01), * (p < 0.05), ‘.’ (p < 0.1).

### Boolean implication network construction

A Boolean implication network (BIN) is created by identifying all significant pairwise Boolean implication relationships (BIRs) for GSE125583 datasets (**Fig. 1b**). The Boolean implication network contains the six possible Boolean relationships between genes in the form of a directed graph with nodes as genes and edges as the Boolean relationship between the genes. The nodes in the BIN are genes and the edges correspond to BIRs. Equivalent and Opposite relationships are denoted by undirected edges and the other four types (low => low; high => low; low => high; high => high) of BIRs are denoted by having a directed edge between them. The network of equivalences seems to follow a scale-free trend; however, other asymmetric relations in the network do not follow scale-free properties. The AD dataset GSE125583 was prepared for Boolean analysis by filtering genes that had a reasonable dynamic range of expression values. When the dynamic range of expression values was small, it was difficult to distinguish if the values were all low or all high or there were some high and some low values. Thus, it was determined to be best to ignore them during Boolean analysis. The filtering step was performed by analyzing the fraction of high and low values identified by the StepMiner algorithm ^29^. Any probe set or genes which contained less than 5% of high or low values were dropped from the analysis.

### Clustered Boolean Implication network

Clustering was performed in the Boolean implication network to dramatically reduce the complexity of the network (**Fig. 1b**). A clustered Boolean implication network (CBIN) was created by clustering nodes in the original BIN by following the equivalent BIRs. One approach is to build connected components in an undirected graph of Boolean equivalences. However, because of noise the connected components become internally inconsistent e.g. two genes opposite to each other becomes part of the same connected component. In order to avoid such situation, we need to break the component by removing the weak links. To identify the weakest links, we first computed a minimum spanning tree for the graph and computed Jaccard similarity coefficient for every edge in this tree. Ideally if two members are part of the same cluster they should share as many connections as possible. If they share less than half of their total individual connections (Jaccard similarity coefficient less than 0.5) the edges are dropped from further analysis. Thus, many weak equivalences were dropped using the above algorithm leaving the clusters internally consistent. We removed all edges that have Jaccard similarity coefficient less than 0.5 and built the connected components with the rest. The connected components were used to cluster the BIN which is converted to the nodes of the CBIN. Increasing the Jaccard similarity cut-off will result in more compact and correlated clusters in CBIN. The distribution of cluster sizes was plotted in a log-log scale to observe the characteristic of the Boolean network. To ensure that the cluster sizes exhibit scale-free properties, the Jaccard similarity cut-off is modified such that they are evenly distributed along a straight line on a log-log plot. A new graph was built that connected the individual clusters to each other using Boolean relationships. Genes in each cluster is ranked based on the number of equivalences within the cluster. Link between two clusters (A, B) was established by using the top representative node from A that was connected to most of the member of A and sampling 6 nodes from cluster B and identifying the overwhelming majority of BIRs between the nodes from each cluster. The 6 nodes include the top representative gene (first rank), the gene next to top (second rank), middle (floor(n/2)^th^ rank where n is the cluster size), gene next to middle (floor(n/2) – 1 rank), middle from top half (floor(n/4)^th^ ranked gene), and middle from the top 1/4^th^ (floor(n/8)^th^ ranked gene) representative nodes from cluster B if size of the cluster is greater than 10. If size of the cluster is between 2 to 10, top two and middle one is picked to test the relationship with cluster A. If the size of the cluster is 1, then it is used to test the relationship with cluster A. Testing multiple nodes provides the most common type of relationships found between cluster A and B. We suggest referring the codebase released for additional details.

A CBIN was created using the selected GSE125583 datasets. The edges between the clusters represented the Boolean relationships that are color-coded as follows: orange for low => high, dark blue for low => low, green for high => high, red for high => low, light blue for equivalent and black for opposite.

### Composite score for clusters of genes

To compute the score, first the genes present in each cluster were normalized and averaged. Gene expression values were normalized according to a modified Z-score approach centered around StepMiner threshold (formula = (expr – SThr-0.5)/3*stddev). Weighted linear combination of the averages from the clusters of genes was used to create a score for each sample. The sample order is visualized by a color-coded bar plot and a violin+swarm plot (**Fig. 1e**). A noise margin is computed for this composite score which follows the same linearly weighted combined score on 2-fold change (+/- 0.5 around StepMiner threshold).

### Training Boolean models to predict AD states

The clusters of genes are selected based on how strongly they predict AD in the training datasets. The genes in these cluster are filtered based on ROC-AUC values (> 0.6 for up regulated and < 0.3 for down regulated) in all three training datasets. The final set of up and down regulated genes are ranked based on T statistic in GSE125583 and top 20 genes were selected for up and down. Weights of up and down regulated genes are 1 and -1 respectively to compute the composite score for the trained AD model.

### RNA-seq library preparation and analysis

Total RNA was isolated from the prefrontal cortex of 8-month-old mice using the RNeasy Mini Kit (Qiagen). RNA quantity and purity were measured by NanoDrop, and integrity was verified using an Agilent TapeStation 4200 system. RNA-seq libraries were constructed from 500 ng of total RNA using the KAPA mRNA HyperPrep Kit (Roche) with Unique Dual-Indexed Adapters (KAPA Biosystems), according to the manufacturer’s protocols. Libraries were PCR-amplified (10 cycles), assessed for quality using TapeStation, and quantified using a Qubit 2.0 fluorometer (Thermo Fisher). Pooled libraries were sequenced on an Illumina NovaSeq 6000 platform using paired-end 100 bp reads at the UCSD Institute for Genomic Medicine (IGM) Core.

### Immunohistochemistry

Mice were anesthetized with isoflurane and transcardially perfused with phosphate-buffered saline (PBS). Brains were fixed in zinc-formalin (Z-fix, Anatech) for 48 hours, followed by cryoprotection in 30% sucrose for 72 hours at 4°C. Coronal sections (30 µm thick) were cut using a sliding freezing microtome (Epredia) and stored at −20°C in cryoprotectant solution. Sections (7–8 per mouse) encompassing anterior to posterior hippocampal regions were selected for staining. After thorough PBS washing (6 × 10 min), sections were incubated with MC1 antibody (1:500; gift from Dr. Peter Davies) in PBS containing 0.4% Triton X-100 for 24 hours at 4°C. Sections were then washed (3 × 15 min PBS), incubated with a fluorescent secondary antibody and DAPI (1:2000) for 1 hour at room temperature, followed by three additional PBS washes. Slides were mounted with Fluoromount-G (Southern Biotech) and imaged using a Keyence BZ-X800 fluorescence microscope.

### Transmission Electron Microscopy (TEM)

Mice were deeply anesthetized and perfused with warm (37°C) Hank’s Balanced Salt Solution (HBSS) containing calcium and magnesium, followed by 2.5% glutaraldehyde and 2% paraformaldehyde in 0.15 M sodium cacodylate buffer using a peristaltic pump as described previously ^48^. Brains were dissected, and hippocampus and cortex were post-fixed in 1% osmium tetroxide (OsO₄) in 0.1 M cacodylate buffer. Tissue blocks were en bloc stained with 2–3% uranyl acetate, dehydrated through an ethanol series, and embedded in resin. Ultrathin sections (50–60 nm) were collected and stained with 2% uranyl acetate and Sato’s lead solution. Images were acquired using a JEOL JEM-1400Plus transmission electron microscope equipped with a Gatan OneView 4k × 4k digital camera ^49^.

### Morphometric analysis

To minimize bias, images were randomly collected and analyzed by three blinded investigators. A total of 25 electron micrographs were evaluated per group (n = 3 animals per group). Vesicle diameter and area were quantified using the line tool in ImageJ (NIH), and vesicle and axon terminal areas were traced using the freehand tool. Vesicle density was calculated by dividing the total vesicle area by the area of the axon terminal as described previously ^50^.

### Statistical Analyses

Gene signature is used to classify sample categories and the performance of the multi-class classification is measured by ROC-AUC (Receiver Operating Characteristics Area Under The Curve) values. A color-coded bar plot is combined with a density or violin+swarm plot to visualize the gene signature-based classification. All statistical tests were performed using R version 3.2.3 (2015-12-10). Standard t-tests were performed using python scipy.stats.ttest_ind package (version 0.19.0) with Welch’s Two Sample t-test (unpaired, unequal variance (equal_var = False), and unequal sample size) parameters. Multiple hypothesis corrections were performed by adjusting *p* values with statsmodels.stats.multitest.multipletests (fdr_bh: Benjamini/Hochberg principles). The results were independently validated with R statistical software (R version 3.6.1; 2019-07-05). Pathway analysis of gene lists was carried out via the Reactome database and algorithm ^51^. Reactome identifies signaling and metabolic molecules and organizes their relations into biological pathways and processes. Kaplan-Meier analysis is performed using lifelines python package version 0.14.6. GraphPad Prism (version 10.5.0) software (San Diego, CA) was used for analysis of electron microscopical and immunohistochemical data. Data were analyzed using unpaired 2-tailed Student *t* test for comparison of 2 groups or 1-way or 2-way ANOVA for >2 groups followed by Sidak’s multiple comparison test if appropriate. For synaptic vesicle area and diameter data, Kolmogorov-Smirnov test was used. All data are presented as mean +SEM. Significance was assumed when *P* was <0.05.

## Results

### Boolean Network Modeling Identifies a Robust and Invariant Alzheimer’s Disease Gene Signature

We analyzed a large-scale bulk RNA-Seq dataset (GSE125583; 70 healthy controls [HC], 219 AD cases) using our Boolean Network Explorer (BoNE) framework to construct a clustered Boolean implication network (“AD-net”, **Fig. 1a–b**). Applying AI/ML-guided refinement across three independent datasets—GSE125583 (fusiform gyrus), GSE15222 (cortex), and GSE118553 (entorhinal cortex)—we identified a core 40-gene Alzheimer’s signature, comprising 20 upregulated and 20 downregulated genes (**Fig. 1c**). Reactome pathway enrichment of this Boolean-derived signature revealed key pathways distinguishing AD from normal aging (**Fig. 1d**). Composite score of the 20 Up and 20 Down genes were used to compute the AD model score. The model score was evaluated on the RNA-Seq dataset (GSE125583; Entorhinal Cortex tissue, 24 healthy controls [HC], 37 AD cases) to check how it differentiates HC with AD samples (**Fig. 1e**). ROC-AUC and two-sided T-test with unequal variance are used to evaluate the performance metric.

We benchmarked this signature against 24 published AD gene sets across all three training datasets. Our Boolean model consistently outperformed these benchmarks in classification accuracy based on average ROC-AUC values (**Fig. 1f–h**), demonstrating superior diagnostic performance and generalizability.

### Boolean AD Signature Validated Across 35 Independent Human Datasets

To further evaluate model robustness, we applied the BoNE signature and the 24 published signatures to 35 independent AD RNA-Seq datasets. The BoNE signature showed the highest mean ROC-AUC values across datasets, outperforming all comparators (**Fig. 2a–d**).

**Figure 2:**
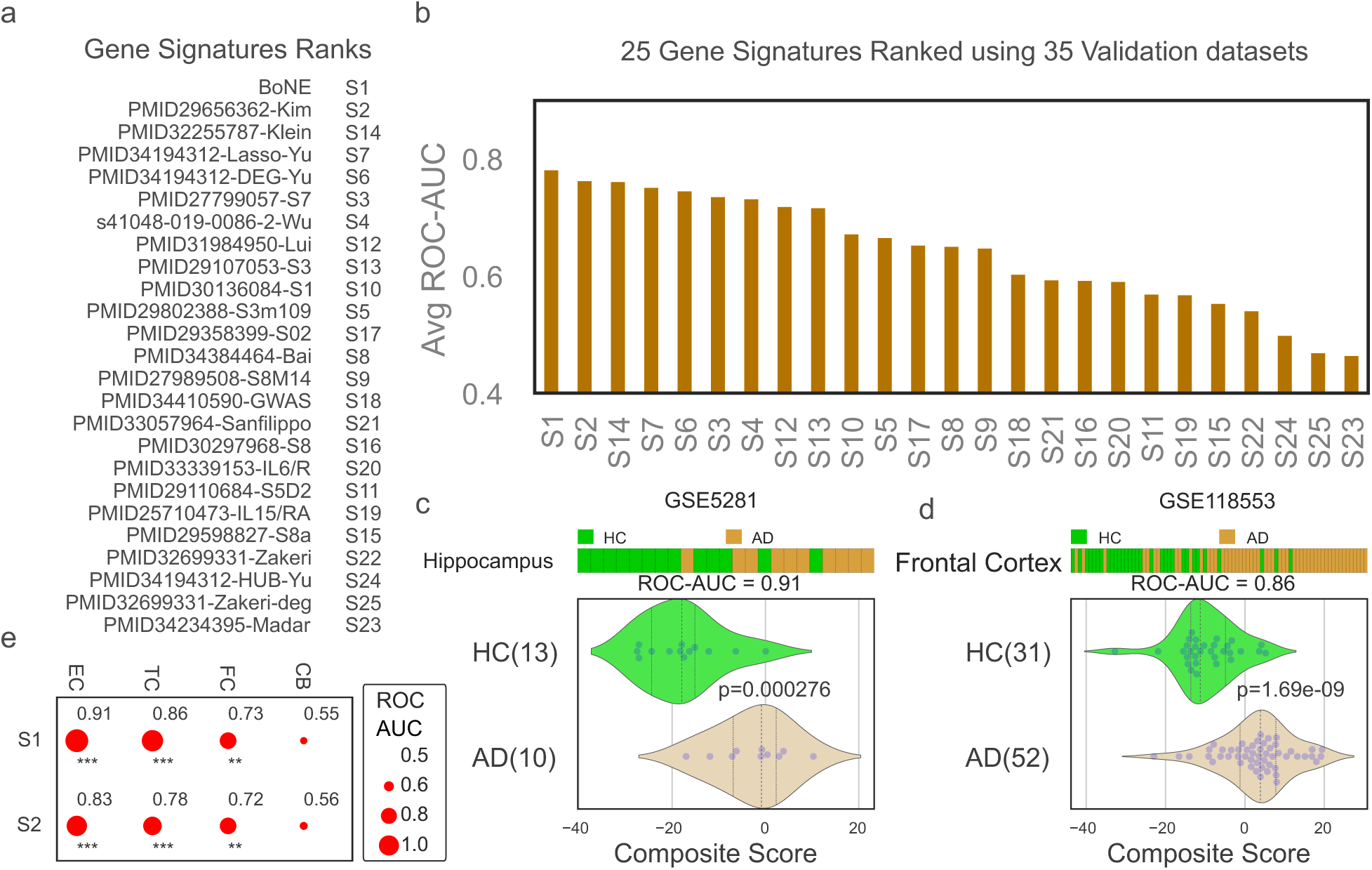
Validation of Boolean model in independent datasets. (**a**) The validation process involved ranking 24 publicly available gene signatures based on their average ROC-AUC values in distinguishing healthy control (HC) and Alzheimer’s disease (AD) samples across 35 independent validation datasets (See Supplementary Data). (**b**) The results are presented in a bar plot, with the x-axis representing the ranked gene signatures and the average ROC-AUC values ordered from highest to lowest. (**c**) The composite score of the Boolean AD models was evaluated, ordering of samples were visualized using bar plot and its distribution in each category is shown in violin plots. The ROC-AUC value and p-value from a two-sided unpaired T-test with unequal variance were computed to assess the Boolean model’s ability to distinguish between 13 HC (colored green) and 10 AD samples (colored orange) from the Hippocampus (GSE5281). (**d**) Bar, violin plot, ROC-AUC, and T-test were performed on another dataset, GSE118553, which included 31 HC and 52 AD samples from the frontal cortex. (**e**) A bubble plot was used to visualize the performance of two signatures, S1 and S2, in different brain regions, including the entorhinal cortex (EC), temporal cortex (TC), frontal cortex (FC), and cerebellum (CB). Bubble size corresponds to ROC-AUC, significance of p values are shown as *** (p < 0.001), ** (p < 0.01), * (p < 0.05), ‘.’ (p < 0.1).

Given that AD pathology typically begins in the entorhinal cortex and hippocampus before spreading to association cortices, we assessed regional BoNE scores in GSE118553. BoNE scores recapitulated the expected topography of disease burden (**Fig. 2e**), reinforcing spatial relevance of the model.

### Boolean Model Enables Discovery of a Murine Model of Asymptomatic AD (AsymAD)

We applied the BoNE signature to five transgenic mouse models of AD (**Fig. 3a**). Ranked sample scores were visualized with WT in green and AD in yellow. BoNE demonstrated high classification accuracy (ROC-AUC: 0.88–1.00) and significant group separation (p < 0.05, unpaired t-test).

**Figure 3:**
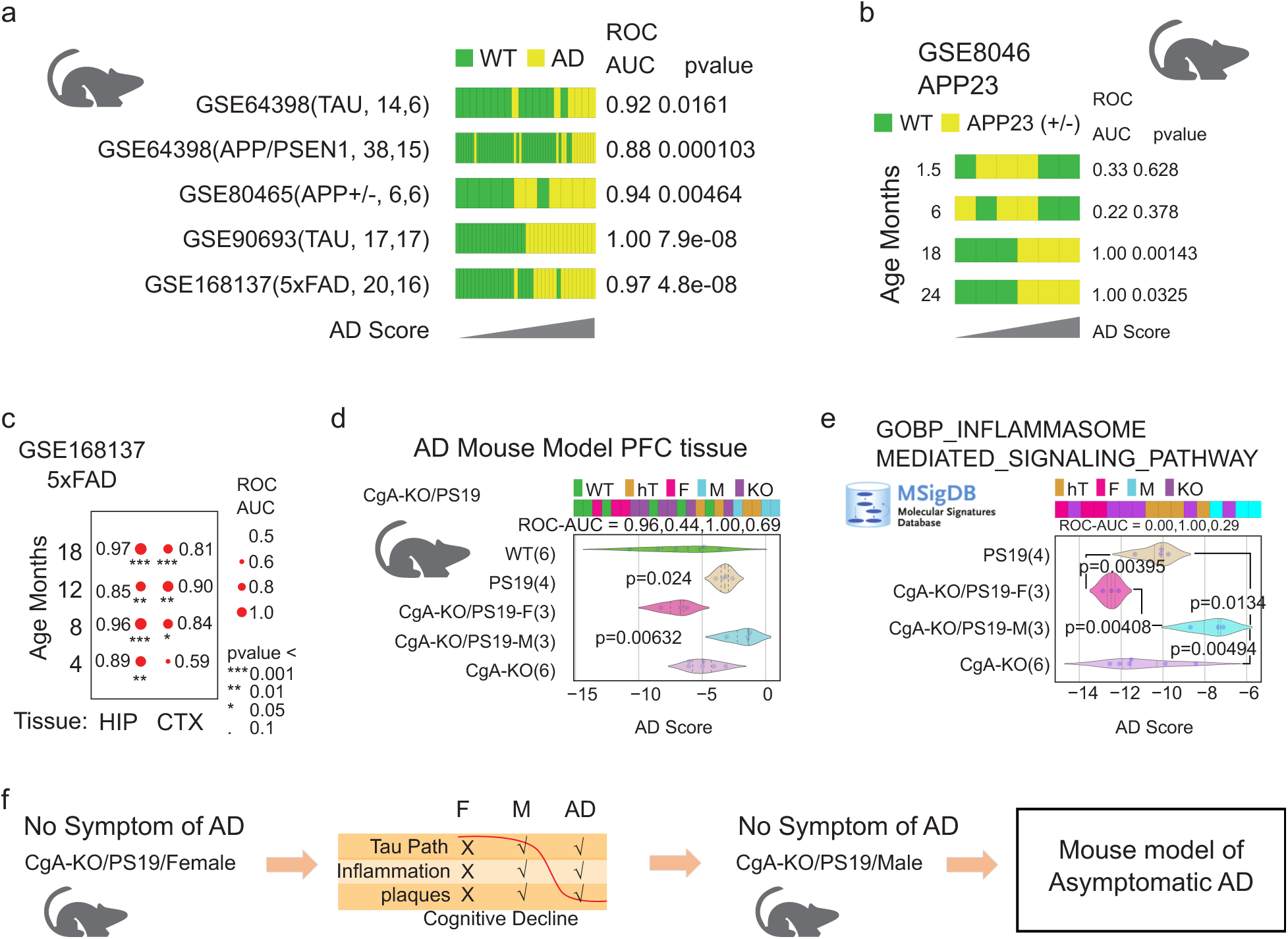
Discovery of Mouse model of AsymAD. (**a**) Boolean AD model derived from human dataset was tested in five different mouse models of AD. Bar plots shows the sample ordering of the composite score from low to high. WT samples are color coded with green and AD samples are color coded with yellow. ROC-AUC and p value from a two-sided unpaired T-test with unequal variance is shown for each dataset. (**b**) WT and APP23 (+/-) AD samples are compared in four different age groups (1.5, 6, 18, 24 months; GSE8046). The results are visualized using bar plots and ROC-AUC and p value from a two-sided unpaired T-test with unequal variance is shown for each dataset. (**c**) WT and 5xFAD AD samples are compared in four different age groups (4, 8, 12, 18 months; GSE168137) and two different tissues (Hippocampus and cortex). The results are visualized in a Bubble plot. Bubble size corresponds to ROC-AUC, significance of p values are shown as *** (p < 0.001), ** (p < 0.01), * (p < 0.05), ‘.’ (p < 0.1). (**d**) Bar, violin plot, ROC-AUC and T-tests were performed using the Boolean AD model composite score on a dataset with PFC samples from five different groups of mice (WT, PS19, CgA-KO/PS19 Female, CgA-KO/PS19 Male, CgA-KO). P values shown are based on two-sided unpaired T-tests with unequal variance comparing WT vs the other four groups of mice. (**e**) Bar, violin plot, ROC-AUC and T-tests were performed using the composite score of genes described in the MSigDB GOBP INFLAMMASOME MEDIATED SIGNALING PATHWAY on a dataset with PFC samples from four different groups of mice (PS19, CgA-KO/PS19 Female, CgA-KO/PS19 Male, CgA-KO). P values shown are based on two-sided unpaired T-tests with unequal variance. (**f**) Boolean AD model discover that CgA-KO/PS19 Male mice PFC tissue have the AD disease states without any symptoms of the AD. Thus, this may represent the world’s first mouse model of the human AsymAD.

In the APP23 model (GSE8046), BoNE scores diverged between WT and AD mice only at 18 and 24 months (Fig. 3b). In the 5xFAD model (GSE168137), the hippocampus—but not cortex—showed robust disease stratification at early stages (**Fig. 3c**).

Most strikingly, in the prefrontal cortex dataset of WT, PS19, CgA-KO, and CgA-KO/PS19 mice (male and female), BoNE scores clustered CgA-KO/PS19 males with PS19, while CgA-KO/PS19 females clustered with WT and CgA-KO (**Fig. 3d**). Despite high BoNE scores (indicative of AD-like molecular pathology), CgA-KO/PS19 males remained behaviorally normal—mirroring the cognitive resilience seen in AsymAD.

Pathway analysis revealed that inflammasome signaling (MSigDB GOBP) was significantly upregulated in PS19 and CgA-KO/PS19 males but suppressed in CgA-KO/PS19 females (**Fig. 3e**). These results identify male CgA-KO/PS19 mice as the first preclinical murine model of AsymAD, characterized by molecular pathology without behavioral deficits.

### Synaptic Vesicle Density Is Preserved in CgA-KO/PS19 Female Mice

Synapse loss and depletion of clear synaptic vesicles (CSVs, ∼40–50 nm) and dense core vesicles (DCVs, ∼80–120 nm) are early hallmarks of AD and correlate with cognitive decline ^52–55^. Electron microscopy of cortical synapses revealed dense CSV clusters in WT males and females (**Fig. 4a-b**). In contrast, PS19 mice of both sexes showed reduced vesicle density (**Fig. 4c-d**), as did CgA-KO/PS19 males (**Fig. 4e**). Remarkably, CgA-KO/PS19 females retained abundant CSVs, resembling WT mice (**Fig. 4f**). This preservation of presynaptic vesicle architecture may underlie their cognitive resilience.

**Figure 4:**
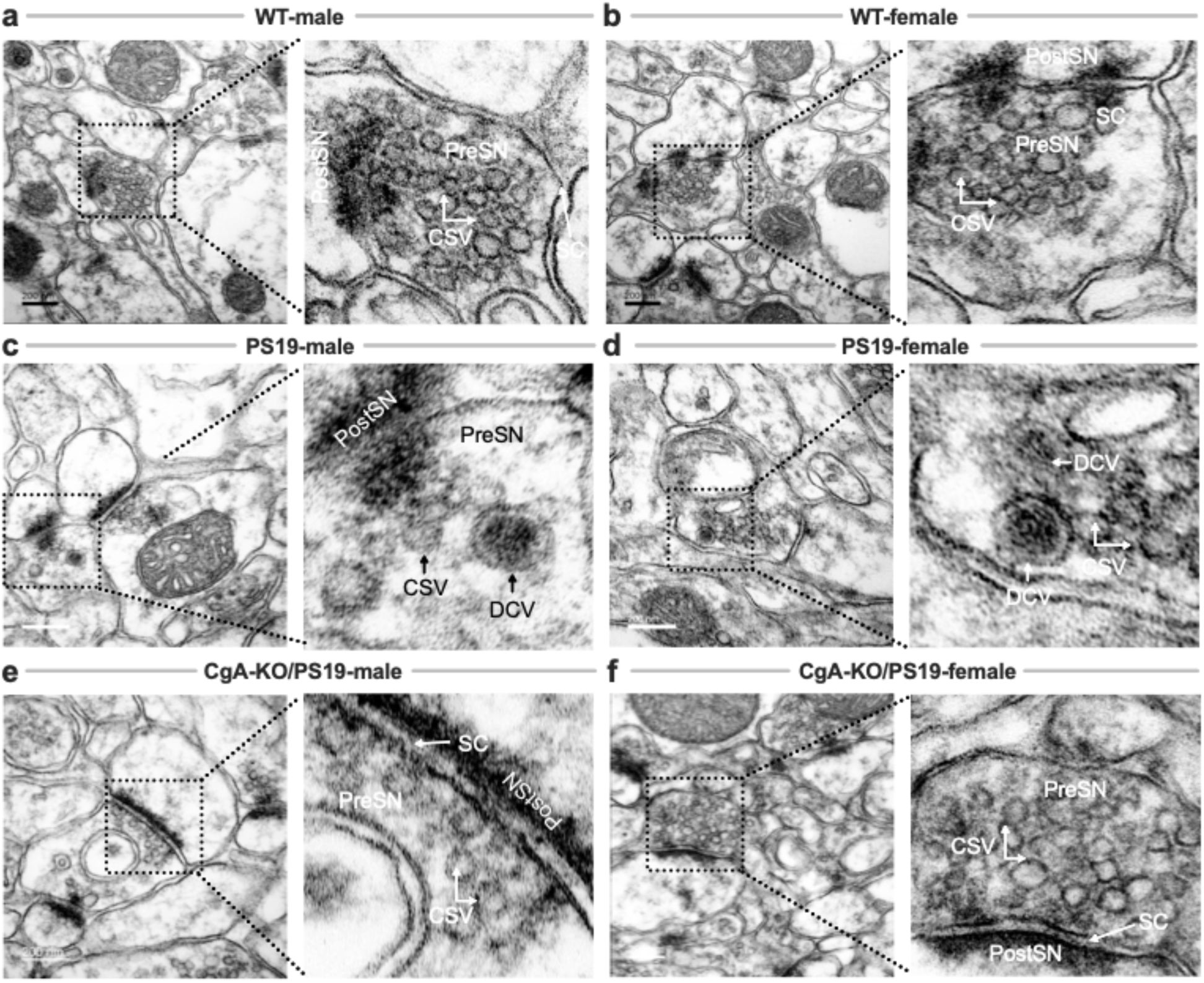
Ultrastructural Analysis of Synaptic Vesicles in the pre-frontal cortices of WT, PS19, and CgA-KO/PS19 Mice. Representative transmission electron micrographs of pre-frontal cortical synapses from wild-type (WT), PS19, and CgA-KO/PS19 mice of both sexes. (**a&b**) WT male (**a**) and female (**b**) mice show abundant clear synaptic vesicles (CSVs) within presynaptic terminals (PreSN), indicative of intact synaptic architecture. (**c&d**) PS19 male (**c**) and female (**d**) mice exhibit markedly reduced CSV density and disrupted synaptic organization, consistent with tauopathy-associated synaptic degeneration. (**e**) CgA-KO/PS19 male mice display a similarly diminished CSV pool as PS19 mice, suggesting persistent synaptic impairment. (**f**) In contrast, CgA-KO/PS19 female mice show restoration of CSV density comparable to WT, indicating preserved synaptic integrity. Scale bars, 200 nm. PreSN: presynaptic terminal; PostSN: postsynaptic terminal; CSV: clear synaptic vesicle; DCV: dense-core vesicle; SC: synaptic cleft.

### Sex-Dependent Accumulation of Neurofibrillary Tangles in the cortex

Multiple studies have shown that women—especially in late age—exhibit greater cortical NFT accumulation than men ^56–59^. Consistent with this, NFTs were absent in WT mice (**Fig. 5a-b**) but abundant in PS19 males and females (**Fig. 5c-d**). CgA-KO/PS19 males mirrored the PS19 pathology (**Fig. 5e**), while CgA-KO/PS19 females were largely devoid of cortical NFTs (**Fig. 5f**). This sex-specific resilience aligns with human AsymAD patterns and underscores a protective role of CgA deletion in females.

**Figure 5:**
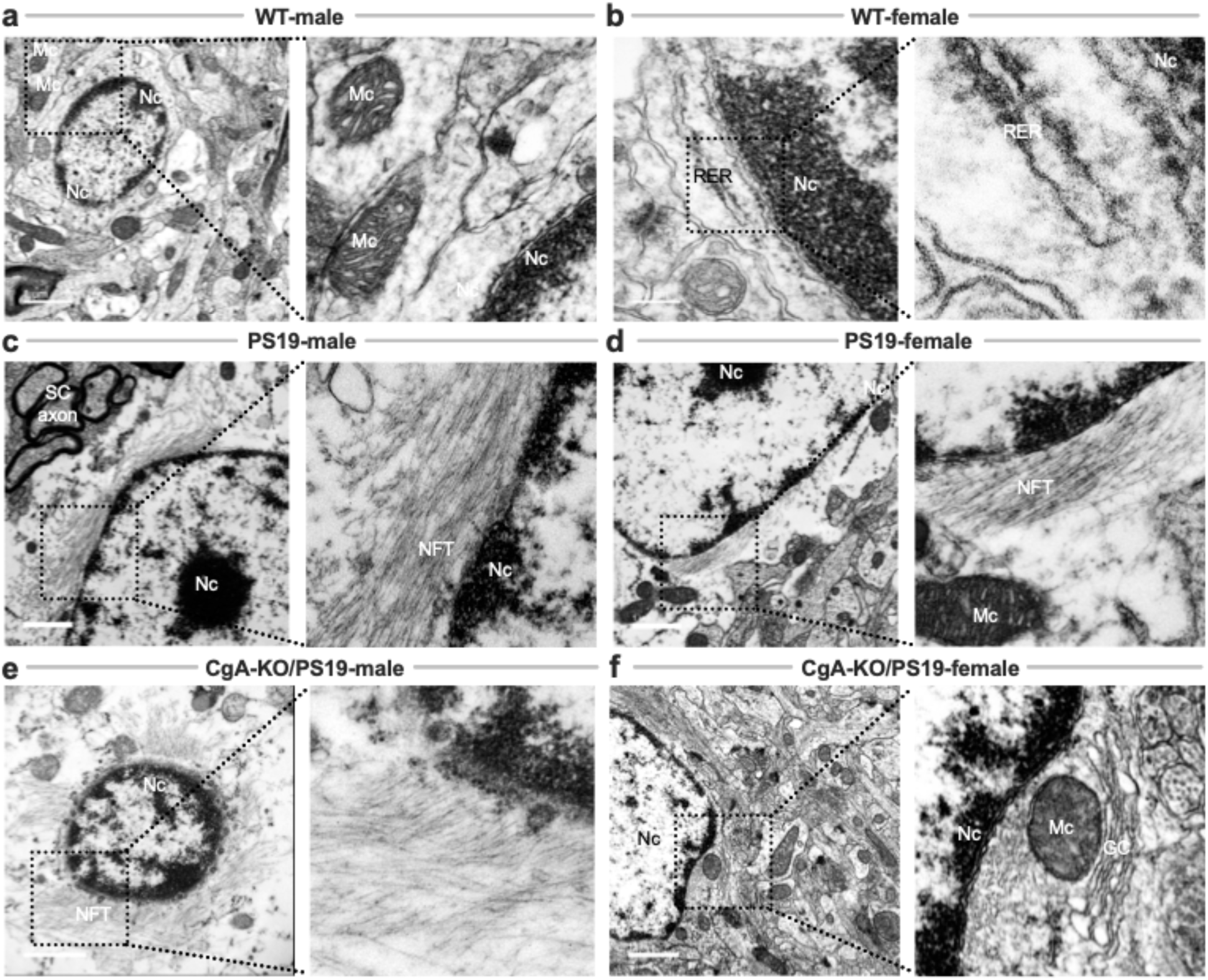
Neurofibrillary Tangle (NFT) Accumulation in Dendritic Regions of PS19 and CgA-KO/PS19 Mice. Transmission electron microscopy (TEM) images of pre-frontal cortical neurons from WT, PS19, and CgA-KO/PS19 mice highlight the presence or absence of neurofibrillary tangles (NFTs) in dendritic regions. (**a&b**) WT male (**a**) and female (**b**) pre-frontal cortices show healthy neuronal morphology with prominent nuclei (Nc), mitochondria (Mc), and intact rough endoplasmic reticulum (RER), with no evidence of NFT accumulation. (**c&d**) PS19 male (**c**) and female (**d**) pre-frontal cortices exhibit extensive NFTs surrounding neuronal nuclei, consistent with advanced tauopathy. (**e**) CgA-KO/PS19 male mice retain dense NFT deposition in dendritic regions, similar to PS19. (F) In contrast, CgA-KO/PS19 female mice lack NFTs and display preserved neuronal ultrastructure comparable to WT controls. Scale bars: A, C, D, E, F = 1 µm; B = 200. nm. Nc: nucleus; Mc: mitochondria; RER: rough endoplasmic reticulum; NFT: neurofibrillary tangle; SC: synaptic cleft; GC: Golgi complex.

### NFT Accumulation in Axons and Axon Terminals Recapitulates Dendritic Trends

Emerging data suggest that NFTs accumulate not only in dendrites but also in axons and synaptic terminals in a sex-dependent manner ^56–59^. PET imaging and postmortem studies consistently show greater cortical tau accumulation in women despite equivalent amyloid loads. This suggests female-specific amplification of tau pathology via axonal mechanisms.

EM revealed NFTs in the axons of PS19 males and females (**Fig. 6c-d**) and in CgA-KO/PS19 males (**Fig. 6e**), but not in WT or CgA-KO/PS19 females (**Fig. 6a-b**, **6f**). These results suggest that tau pathology in axons mirrors dendritic trends and supports the sexually dimorphic regulation of tau aggregation in AsymAD.

**Figure 6:**
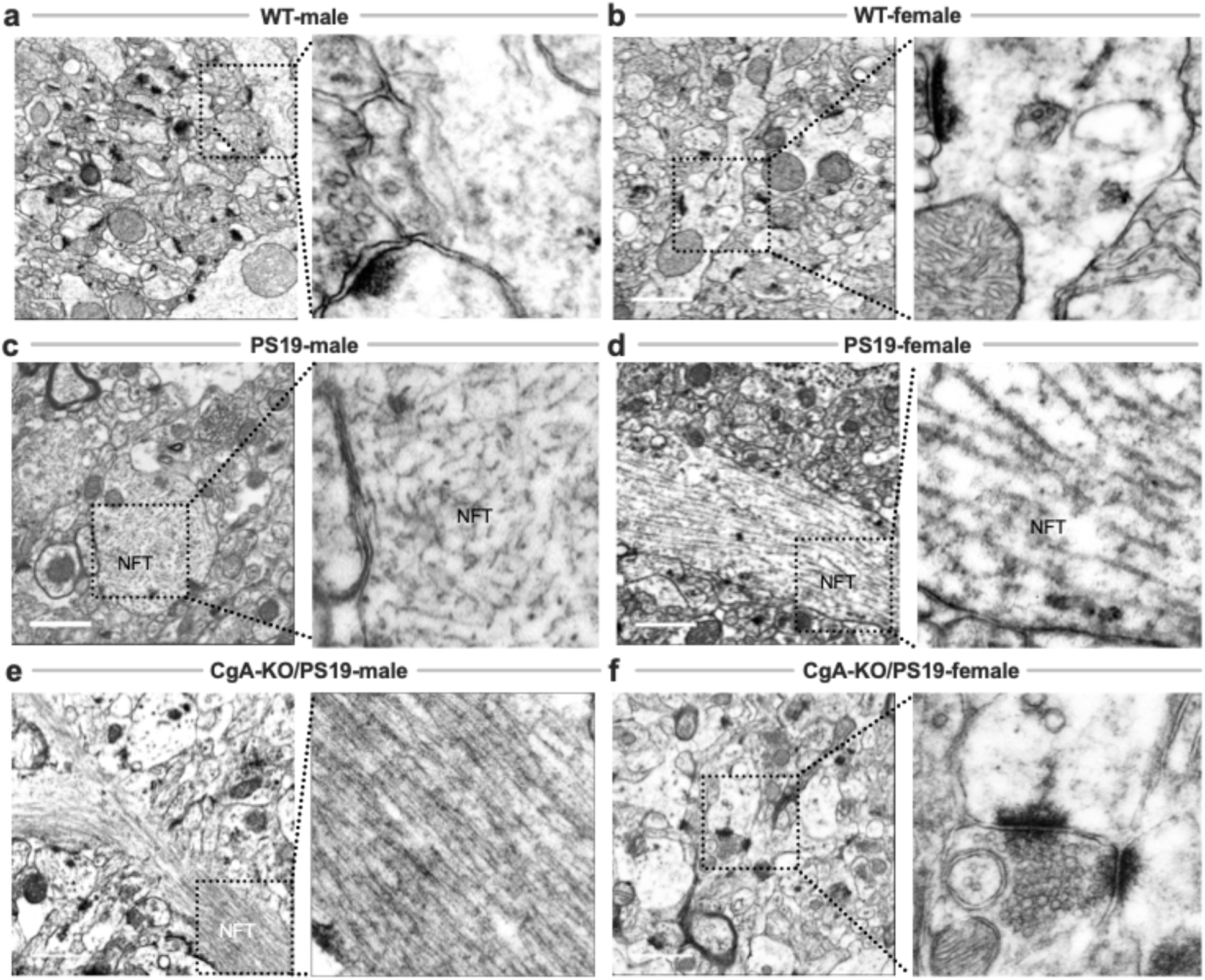
Neurofibrillary Tangles (NFTs) in Axons and Axon Terminals of PS19 and CgA-KO/PS19 Mice. Transmission electron microscopy (TEM) images of pre-frontal cortical axons and axon terminals from WT, PS19, and CgA-KO/PS19 mice highlight NFT distribution in tauopathy. (**a&b**) WT male (**a**) and female (**b**) pre-frontal cortices show normal axonal ultrastructure with no evidence of NFT accumulation. (**c&d**) PS19 male (**c**) and female (**d**) mice exhibit dense filamentous NFTs within axons and axon terminals, characteristic of advanced tau pathology. (**e**) CgA-KO/PS19 male mice show NFT deposition comparable to PS19, indicating sustained tauopathy. (**f**) In contrast, CgA-KO/PS19 female mice lack detectable NFTs and exhibit preserved axonal morphology, mirroring WT. Scale bars: 1 µm. NFT: neurofibrillary tangle.

### Reduced misfolded Tau aggregation in CgA-KO/PS19 Female Mice

Misfolded Tau aggregation was abundant in PS19 males and females (**Fig. 7a**, **7c**) and in CgA-KO/PS19 males (**Fig. 7b**). However, CgA-KO/PS19 females displayed a marked reduction in misfolded Tau aggregation (**Fig. 7d**). Quantitative analysis confirmed a statistically significant reduction in misfolded-tau only in females (**Fig. 7e**), consistent with previous findings that CgA deficiency attenuates tau pathology via sex-dependent mechanisms ^28^.

**Figure 7:**
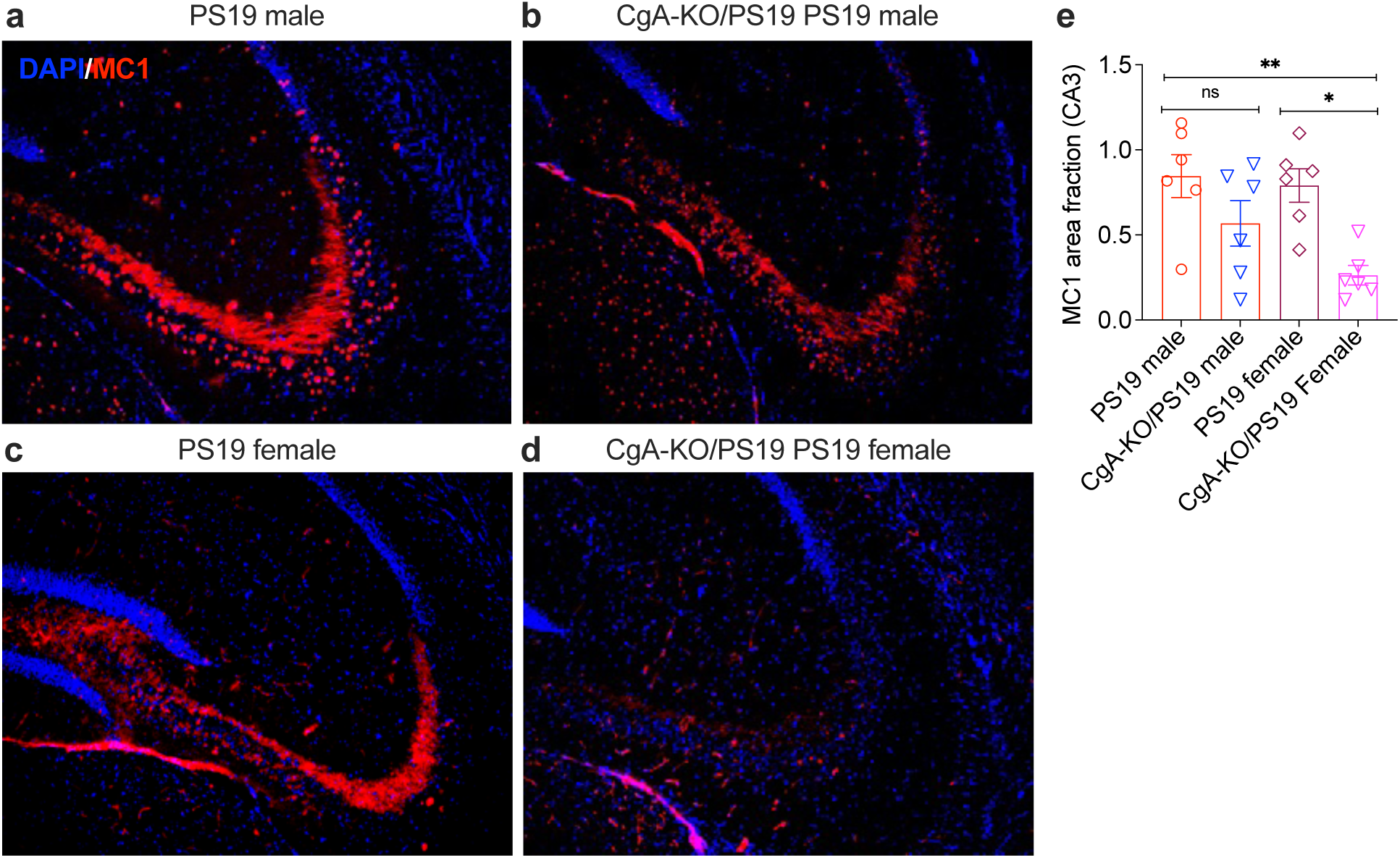
CgA deficiency reduces misfolded Tau aggregation in female PS19 mice. Representative hippocampal sections from (**a**) PS19 male, (**b**) CgA-KO/PS19 male, (**c**) PS19 female, and (**d**) CgA-KO/PS19 female mice were stained for misfolded Tau aggregation using the MC1 antibody (red) and counterstained with DAPI (blue). MC1 immunoreactivity is prominent in both PS19 males and females. While CgA-KO/PS19 males show comparable MC1 staining to PS19 males, CgA-KO/PS19 females exhibit markedly reduced MC1 signal. (**e**) Quantification of MC1-positive area fraction in the CA3 region of the hippocampus shows a significant reduction in Tau oligomer burden in CgA-KO/PS19 females compared to PS19 females (*p < 0.05), with no significant difference between PS19 and CgA-KO/PS19 males (ns). Data are presented as mean ± SEM; *p < 0.05, **p < 0.01 by one-way ANOVA with post hoc test.

## Discussion

We report the first validated murine model of Asymptomatic Alzheimer’s Disease (AsymAD), integrating systems-level Boolean network modeling with multimodal behavioral, molecular, and neuropathological interrogation. AsymAD is a preclinical phase of Alzheimer’s disease (AD) observed in ∼20–30% of cognitively intact elderly individuals who nonetheless harbor substantial amyloid and Tau pathology at autopsy ^5–7^. Despite its clinical relevance, the molecular and cellular mechanisms conferring cognitive resilience in AsymAD remain poorly understood ^13–16^.

Using Boolean Network Explorer (BoNE), we trained a logic-based implication network on large-scale transcriptomic datasets from multiple human cortical regions to identify a core invariant AD signature. This 40-gene signature, comprising 20 upregulated and 20 downregulated genes, was identified by training three independent datasets and shown to outperform 24 published AD gene sets across 35 diverse bulk and single-cell RNA-seq validation datasets. When applied to our murine model, BoNE revealed that male Chromogranin A–deficient PS19 mice (CgA-KO/PS19) exhibit robust AD-like transcriptomic signatures in the hippocampus and entorhinal cortex yet retain intact cognitive function—recapitulating the clinicopathologic dissociation observed in human AsymAD.

This dissociation offers a powerful *in vivo* platform to probe resilience before the onset of symptomatic neurodegeneration. Importantly, the Boolean modeling approach provides a distinct advantage over traditional differential expression methods by capturing stable, context-invariant gene relationships that reflect disease logic rather than noise or cohort-specific variance. This systems framework enables the derivation of testable hypotheses, moving beyond correlation to uncover mechanistic drivers of disease and protection.

A particularly striking feature of the CgA-KO/PS19 model is the emergence of sex-specific resilience. Male mice exhibited preserved spatial learning and memory despite elevated Tau transcripts and AD-associated gene expression. Female CgA-KO/PS19 mice demonstrated an even more protective phenotype, with attenuated Tau phosphorylation, absence of neurofibrillary tangles (NFTs), preserved synaptic ultrastructure, and reduced inflammatory gene signatures. These findings reinforce accumulating evidence that biological sex is a critical modulator of neurodegenerative susceptibility ^60–63^. Hormonal signaling (e.g., estrogen receptor pathways), sex chromosome dosage, and immune regulatory networks may differentially shape neuronal and glial responses to pathology ^64–66^. Our results suggest that resilience may involve distinct transcriptomic and structural adaptations, and future work should dissect the molecular underpinnings of this sex divergence.

Our study further highlights Chromogranin A (CgA), a neuroendocrine secretory granin, as a modifiable determinant of disease vulnerability. CgA is elevated in the CSF of AD patients ^67^ and colocalizes with NFTs in postmortem brains ^22–24^. We previously showed that CgA deletion in PS19 mice attenuates Tau pathology and improves survival ^28^. Here, we demonstrate that CgA deficiency preserves cognition despite persistent AD-like gene signatures, indicating that CgA may actively drive neurodegeneration through non-transcriptomic mechanisms. CgA and its cleavage products—such as Catestatin— are implicated in catecholaminergic modulation ^66, 67^, neuroimmune signaling ^28, 68, 69^, and synaptic plasticity ^70–72^. These multifunctional roles position CgA as a central integrator of neuroimmune and neuroendocrine crosstalk in AD, and a candidate for therapeutic targeting.

## Limitations and Future Directions

While our findings establish a novel and tractable model of AsymAD, several limitations warrant consideration. First, our transcriptomic and neuropathological analyses were focused on the hippocampus and prefrontal cortex, excluding other regions implicated in early AD, such as the anterior cingulate, basal forebrain, and locus coeruleus. Second, although we observed robust sex differences, the precise contributions of estrogen, X-linked gene expression, and chromatin accessibility remain to be determined. Third, our behavioral data are cross-sectional; longitudinal studies are needed to assess whether the resilience phenotype is stable or transient, and how it evolves with age and pathology. Finally, while Boolean network modeling identifies disease-relevant signatures, future integration of proteomics, metabolomics, electrophysiology, and glial functional states will be critical to bridge transcriptomic logic with neural circuit resilience and therapeutic translation.

## Conclusion

Collectively, this work establishes a robust, experimentally validated framework for dissecting resilience in AD. By fusing Boolean logic modeling with a novel CgA-KO/PS19 mouse model, we define a systems-level blueprint for AsymAD and uncover sex-specific neuroprotective signatures. Chromogranin A emerges as both a marker and modulator of disease susceptibility, with implications for therapeutic targeting in early-stage and preclinical AD. This scalable platform enables mechanistic dissection of resilience circuits and positions the field to advance personalized, preventive interventions in neurodegenerative disease.

## Authors’ contributions

Conceptualization: D.S., S.K.M.

Methodology: D.S., S.T., S.K.M.

Investigation: D.S., S.K.M., S.T., S.J., S.K.

Visualization: D.S., S.K.M.

Funding acquisition: S.K.M, D.S

Project administration: S.K.M., D.S.

Supervision: S.K.M., D.S.

Writing – original draft: D.S., S.K.M., S.T

Writing – review & editing: S.K.M., D.S., S.T., S.J., S.K., S.C.S., B.P.H.

## Data sharing statement

All data are available in the main text or the supplementary materials. The codes are available at https://github.com/sahoo00/ADnet.

## Acknowledgements

This work was supported by the National Institutes for Health (NIH) grant R01-AI155696 (to DS), R21 AG072487-01 (to SKM, Gourisankar Ghosh & Xu Chen), R21 AG078635-01A1 (to SKM and Gourisankar Ghosh) and VA Merit Review Grant I01 BX003934 (to SKM). Other support includes VA Merit BX003671 and VA RCS BX006318 to BPH. CDMRP AL210059 (ALSTIA) to BPH; CDMRP AL230115 (ALSTDA) to BPH, UCSD GTI 2039592 to BPH. Other sources of support include: R01-GM138385 (to DS) and UG3TR003355 (to DS). We also acknowledge support of this work by the Wu Tsai Human Performance Alliance (WTHPA) and the Joe and Clara Tsai Foundation. The research was supported in part by NIDDK grant P30 DK120515 in the form of SDDRC core services.

